# Mate choice enhances post-zygotic barriers to gene flow via ancestry bundling

**DOI:** 10.1101/2021.09.02.458713

**Authors:** Pavitra Muralidhar, Graham Coop, Carl Veller

## Abstract

Hybridization and subsequent genetic introgression are now known to be common features of the histories of many species, including our own. Following hybridization, selection often purges introgressed DNA genome-wide. While mate choice can prevent hybridization in the first place, it is also known to play an important role in post-zygotic selection against hybrids, and thus the purging of introgressed DNA. However, this role is usually thought of as a direct one: a mating preference for conspecifics reduces the sexual fitness of hybrids, reducing the transmission of introgressed ancestry. Here, we explore a second, indirect role of mate choice as a barrier to gene flow. Under assortative mating, parents covary in their ancestry, causing ancestry to be “bundled” in their offspring and later generations. This bundling effect increases ancestry variance in the population, enhancing the efficiency with which post-zygotic selection purges introgressed DNA. Using whole-genome simulations, we show that the bundling effect can comprise a substantial portion of mate choice’s overall effect as a post-zygotic barrier to gene flow, and that it is driven by ancestry covariances between and within maternally and paternally inherited genomes. We derive a simple method for estimating the impact of the bundling effect from standard measures of assortative mating. Applying this method to data from a diverse set of hybrid zones, we find that the bundling effect increases the purging of introgressed DNA by between 1.2-fold (in a baboon system with weak assortative mating) and 14-fold (in a swordtail system with strong assortative mating). Thus, the bundling effect of mate choice contributes substantially to the genetic isolation of species.

---

The preference for mating with conspecifics is an important barrier to gene flow between species (Coyne & Orr 2004). In this capacity, mate choice can act pre-zygotically, preventing the formation of hybrid offspring, and post-zygotically, with hybrids being unattractive to the majority of potential mates (Mayr 1942, Servedio & Noor 2003, Price 2008). Following admixture, post-zygotic factors can cause deleterious effects in hybrids, leading to selection against introgressed DNA. Alongside mate choice, these factors include incompatibility of genetic variants from the two parent species (Dobzhansky 1937, Muller 1942, Orr & Turelli 2001), maladaptation of introgressed alleles to the recipient species’ ecology (Schluter & Conte 2009), and higher genetic load in the donor species (Harris & Nielsen 2016, Juric et al. 2016). Recent evidence suggests that the deleterious effect of post-zygotic factors can often be spread across a large number of genomic loci (Schumer et al. 2014, Juric et al. 2016, Aeschbacher et al. 2017) [e.g., ∼1,000 loci for Neanderthal-human introgression (Juric et al. 2016)].

When introgressed ancestry is deleterious at many loci throughout the genome, the rate at which it is purged by selection is proportional to the variance across individuals in how much introgressed DNA they carry (Fisher 1930, Harris & Nielsen 2016, Veller et al. 2021, Methods). In light of this, we reasoned that, while mate choice contributes directly to the purging of introgressed DNA in a given generation—via the reduced sexual fitness of hybrids—it can also contribute indirectly to purging in the next and later generations, by altering how introgressed DNA is packaged among offspring. Specifically, positive assortative mating “bundles” like-with-like ancestry in the formation of offspring, increasing population-wide ancestry variance in offspring and later generations (Crow & Felsenstein 1968, Goldberg et al. 2020) (Fig. 1). This increased ancestry variance enhances the efficiency with which post-zygotic selection of various kinds purges introgressed DNA in these later generations. Therefore, there exist two mechanisms by which mate choice acts as a barrier to gene flow between species: a direct, “sexual selection” mechanism and an indirect, “bundling” mechanism.

**Figure 1.**
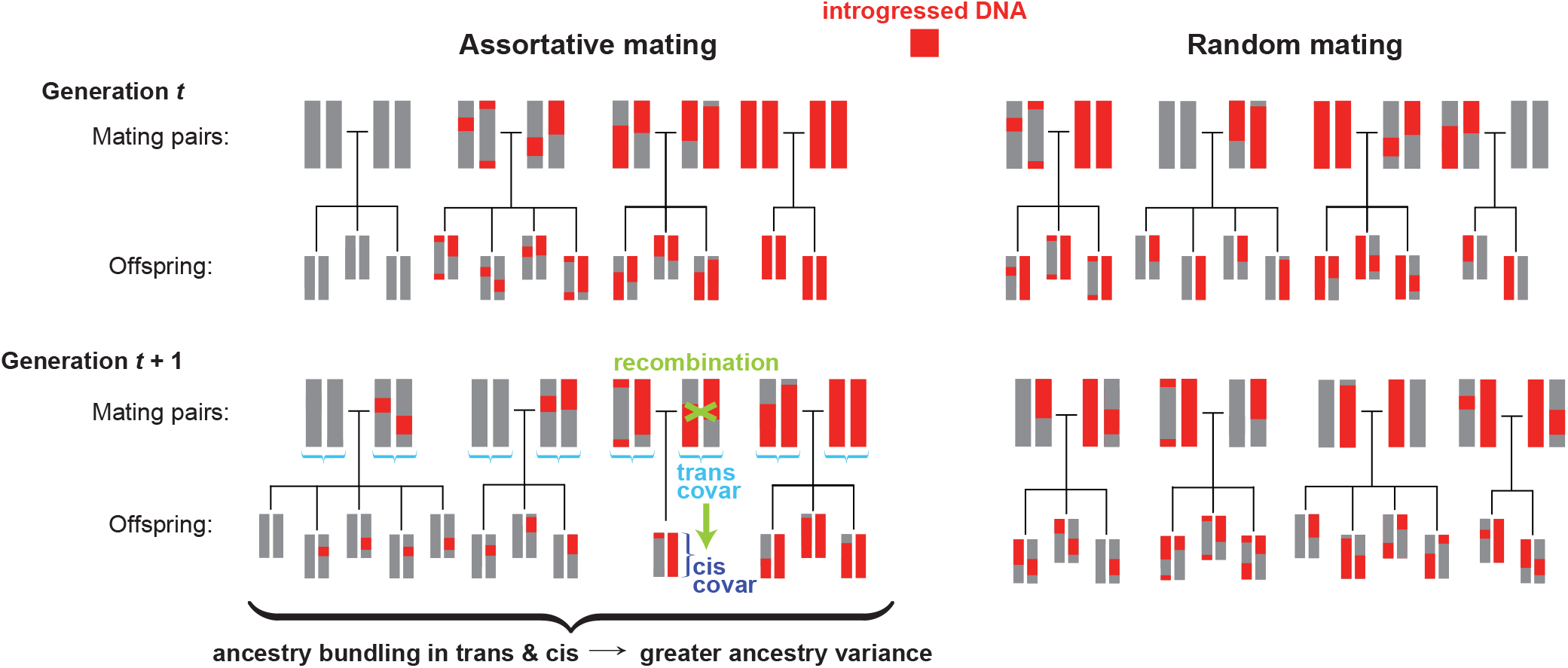
Assortative mating bundles introgressed DNA together, increasing the efficiency with which it is purged by selection. Introgressed alleles initially appear in the population in perfect linkage disequilibrium (cis covariance) with one another; recombination subsequently breaks down these cis covariances over time. A mating preference for conspecifics generates positive ancestry correlations between mating pairs, so that offspring inherit maternal and paternal genomes that covary in their proportions of introgressed DNA—i.e., that covary “in trans”. These “trans covariances” are subsequently converted to new cis covariances, as covarying maternal and paternal genomes recombine into the same gametes. The result is that, relative to random mating, assortative mating causes introgressed DNA to become more densely concentrated in a smaller number of individuals. This “bundling” effect increases the variance across individuals in how much deleterious introgressed DNA they carry, and therefore increases the rate at which introgressed DNA is purged by selection.

To study the contribution of these two mechanisms to the genetic isolation of species, we considered a model in which a recipient and donor species experience a single pulse of admixture, resulting in a fraction of donor DNA admixing into the recipient species’ gene pool. Introgressed alleles at many loci are assumed to reduce viability in the recipient species, such that an individual with introgressed genomic fraction *I* has relative viability fitness 1 − *IS*, where *S* is the strength of viability selection against introgressed ancestry. (This situation could arise, for example, if the donor species is poorly adapted to the recipient species’ local environment.) We initially assume that females exercise mate choice according to a fixed relative preference model (Seger 1985) based on ancestry, with a female preferring to mate with males from the species that matches her majority ancestry. Specifically, if a female’s introgressed fraction is *I*_*f*_, then her probability of mating with a given male of introgressed fraction *I*_*m*_ is proportional to

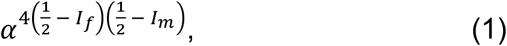

where *α* > 1 quantifies the overall strength of mate choice in the system. Thus, if a female is of 100% recipient-species ancestry (*I*_*f*_ = 0), she prefers males of 100% recipient-species ancestry (*I*_*m*_ = 0) over fully hybrid males (*I*_*m*_ = 1/2) by a factor of *α*, and over 100% donor-species males (*I*_*m*_ = 1) by a factor of *α*^2^. In contrast, fully hybrid females (*I*_*f*_ = 1/2) are indiscriminate in mate choice. We consider alternative specifications of mate choice later.

We performed whole-genome simulations of this model (Haller & Messer 2019), and observed rapid purging of introgressed ancestry following the initial admixture pulse (Fig. 2). This purging is due to (i) viability selection, (ii) sexual selection induced by the direct effect of mate choice, and (iii) the enhancement of (i) and (ii) by the bundling effect of mate choice. To isolate the contributions of mate choice’s sexual selection and bundling effects, we used simulation experiments to artificially eliminate the bundling effect while preserving the fitness consequences of mate choice for males. Each generation, we calculated the sexual fitness of every adult male under the model of mate choice described above (averaged over the population of adult females), and reassigned these sexual fitnesses to viability fitnesses that took effect in an additional round of viability selection. Mating pairs were then formed at random among surviving males and females. This procedure preserves the sexual selection effect of mate choice, since attractive males still enjoy the same higher fitness, but it eliminates the bundling effect of mate choice because the offspring generation is produced by random mating.

**Figure 2.**
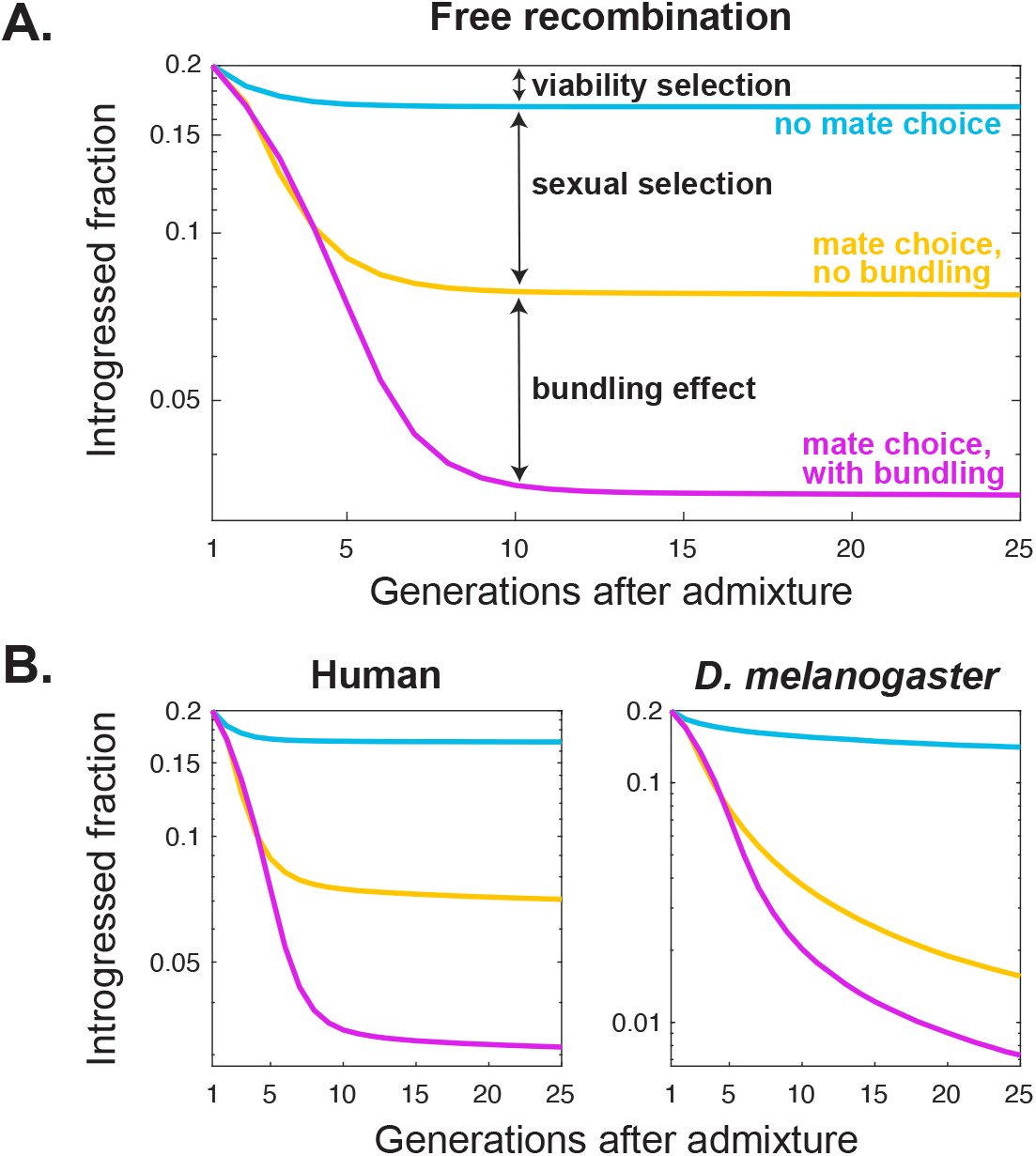
The bundling effect of mate choice is an important contributor to the genetic isolation of species. A. In our model, introgressed ancestry is purged as a result of viability selection and sexual selection, and their enhancement by the bundling effect of mate choice. These three effects can be distinguished using simulation experiments that artificially preserve the sexual selection induced by mate choice but remove its bundling effect. In the absence of the bundling effect, the purging of introgressed ancestry (yellow line) is substantially slower and less profound than in the presence of the bundling effect (purple line). The simulations here assume that all loci are unlinked (*r* = 1/2), *α* = 4, *S* = 0.1, and an initial admixture proportion of 20%. The y-axis is log-scaled, so that trajectory slopes represent rates of purging. **B**. The bundling effect contributes substantially to mate choice’s overall effect in cases of realistic recombination processes. Its contribution is especially large in the case of humans, a high-recombination species.

We found that, in simulations with this unbundling procedure, substantially less introgressed DNA was purged than in simulations with unmanipulated mate choice. Consider the case displayed in Fig. 2A, where all loci are unlinked. In the absence of mate choice, the introgressed fraction would be reduced by viability selection alone from 20% to 16% after 25 generations (Fig. 2A). In the presence of mate choice, the introgressed fraction is in fact reduced to just 3%, so that the overall effect of mate choice is an additional 13 percentage points of purging. However, if we remove the bundling effect of mate choice, the additional purging across 25 generations is just 8.5 percentage points (Fig. 2A). Therefore, in this case, the bundling effect accounts for more than one third of mate choice’s overall effect. Put differently, viability selection would need to be 6.8-fold stronger to match the amount of purging after 25 generations under full mate choice, but only 4.5-fold stronger if the bundling effect of mate choice is removed (Fig. S1). Thus, bundling amplifies mate choice’s effect on the strength of selection against introgressed DNA by ∼50% in this case. As expected, the contribution of bundling increases with the strength of assortative mating (Fig. S2).

The increase in the rate of purging due to the bundling effect derives from increased ancestry variance in the population caused by assortative mating (Fig. 3A). In our unbundling simulations, which impose random mating, the population ancestry variance—normalized for the overall proportion of introgressed ancestry—matched that observed in simulations without mate choice (Fig. 3A). In contrast, when the bundling effect of mate choice is preserved, the ancestry variance substantially exceeds this random-mating expectation (Fig. 3A).

**Figure 3.**
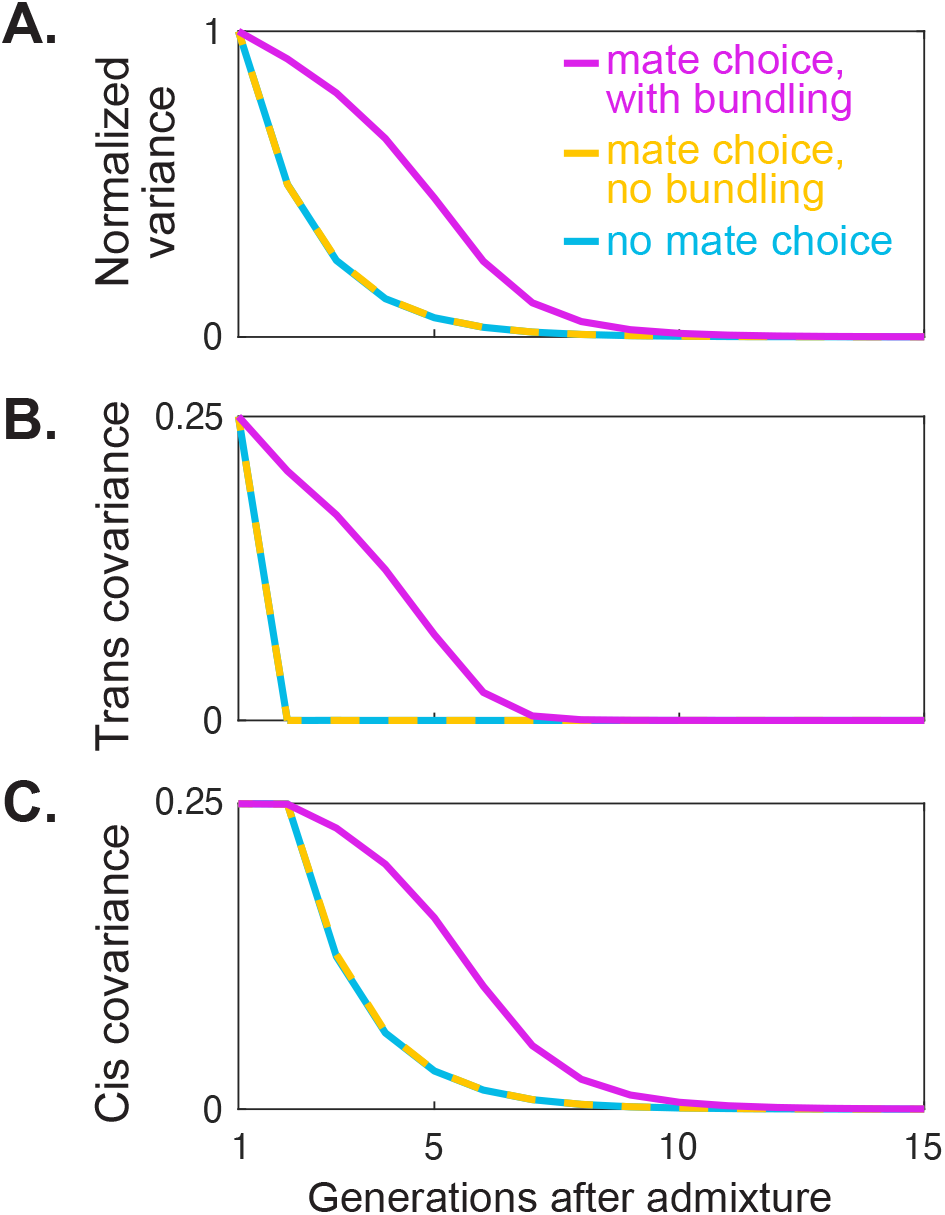
The bundling effect of mate choice increases ancestry variance by generating ancestry covariance between and within maternally and paternally inherited genomes. Evolution of the population’s overall ancestry variance, trans-covariance, and cis-covariance, under the three scenarios of no mate choice, full mate choice, and mate choice with its bundling effect removed. Since introgressed ancestry is purged at different rates in the three cases, and since the overall introgressed fraction *I*_*t*_ influences the range of possible ancestry variances and covariances, we normalize each variance and covariance trajectory by the variance expected from the introgressed fraction alone, absent any linkage disequilibria: *I*_*t*_(1 – *I*_*t*_). **A**. Recombination rapidly breaks down the initial ancestry variance in the population. The decay of the variance is, however, substantially slower with mate choice (purple line) than without mate choice (blue line). This increased variance is due to mate choice’s bundling effect: the normalized trajectory under mate choice with its bundling effect removed is the same as that under no mate choice. **B**. One component of the increased variance caused by the bundling effect is due to ancestry covariances across maternally and paternally inherited genomes, which arise because mate choice causes mating pairs to have correlated ancestries. Without mate choice, or with its bundling effect removed, the trans covariance is zero (except in the first generation where individuals are all of one species or the other). **C**. The second component of the bundling effect is its effect on ancestry covariances within haploid genomes. Cis covariances are initially large in all scenarios (since introgressed alleles appear in the population in perfect linkage disequilibrium), but they are rapidly broken down over time by recombination. Mate choice decelerates this decay, by generating trans covariances which recombination converts into cis covariances. Parameters as in Fig. 2A.

The results described above are for a highly stylized genome, where all loci are unlinked. To see if the bundling effect of mate choice can be important in more realistic genomes, we repeated our simulations using linkage maps for a high-recombination species [humans (Kong et al. 2010)] and a low-recombination species [*Drosophila melanogaster* (Comeron et al. 2012)] (Methods). We found that the bundling effect constitutes a large proportion of mate choice’s overall effect in the human case, and a smaller proportion in *D. melanogaster* (Fig. 2B). This implies that recombination plays an important role in the bundling effect, which in turn highlights a role for linkage disequilibria among introgressed alleles.

We formalized this intuition by decomposing the overall population ancestry variance into three components (Methods): (1) a small component due to heterozygosity at different loci; (2) a component due to ancestry covariance between individuals’ maternally and paternally inherited genomes (trans-linkage disequilibrium, or trans-LD; Fig 1); (3) a component due to ancestry covariance within individuals’ maternally and paternally inherited genomes (cis-linkage disequilibrium, or cis-LD; Fig 1). In the first generation after admixture, all introgressed alleles have been inherited from donor-species parents, and therefore lie in perfect cis-LD with one another. Recombination breaks down this initial cis-LD quickly over subsequent generations (Fig. 3C), reducing ancestry variance and thus slowing the rate of purging (e.g., Harris & Nielsen 2016, Veller et al. 2021). If mating were random, trans-LD would be zero in all generations after the initial admixture event (Fig. 3B), and so the rate of decay of overall ancestry variance would depend only on the changing frequencies of introgressed alleles (component 1) and the reduction in their cis-LD by recombination (component 3).The bundling effect of mate choice impedes the decay of ancestry variance in two ways: by generating trans-LD (component 2; Fig. 3B) and by slowing down the decay of cis-LD (component 3; Fig. 3C).

The “trans channel” of the bundling effect is a direct consequence of mate choice: mating pairs have disproportionately similar ancestry, so offspring inherit maternal and paternal genomes with correlated ancestry—i.e., trans-LD (Fig. 1). While the degree of trans-LD in a given generation, and thus the strength of the trans channel, does not depend on recombination in the previous generation, the “cis channel” is driven largely by recombination converting trans-LD from the previous generation into cis-LD (Fig. 1) [as shown in previous models of mate choice (Kirkpatrick 1982, Zaitlen et al. 2017, Veller et al. 2020)].

Our results thus far have excluded sex chromosomes, which are of particular interest as they are known to be enriched for genes involved in mate choice (Muralidhar 2019) and to show distinct patterns of cis-LD and trans-LD in models of sexual selection (Kirkpatrick & Hall 2004, Veller et al. 2020). Incorporating sex chromosomes into our model, we find that they tend to purge introgressed ancestry at a higher rate than autosomes (Fig. S3), because they do not recombine in the heterogametic sex and therefore maintain longer, more deleterious introgressed linkage blocks than the autosomes. Interestingly, we find that Z chromosomes (in female-heterogametic taxa, such as birds and butterflies) purge more introgressed DNA than X chromosomes (in male-heterogametic taxa, such as mammals and flies) (Fig. S3), with the importance of the bundling effect concomitantly larger for Z chromosomes. This difference can be explained by the fact that, under an even sex ratio, two thirds of Z chromosomes each generation are in males, on whom selection against introgressed ancestry is stronger than in females because of the additional effect of sexual selection. In contrast, under male heterogamety, two thirds of X chromosomes are in females, on whom selection against introgressed ancestry is weaker. Our model assumes female mate choice. While male mate choice is now known to play an important role in many systems (Edward & Chapman 2011), sexual selection tends to be stronger for males than for females (Janicke et al. 2016). Therefore, all else equal, our results show that the influence of mate choice—and concomitantly its bundling effect—on the purging of sex-linked introgressed ancestry is stronger in female-heterogametic species than in male-heterogametic species.

Throughout, we have assumed a simple model in which genome-wide ancestry determines the mating preferences of females, the attractiveness of males, and viability. In a genetically more realistic model, separate loci would underlie these three distinct traits. To check that our results are robust to consideration of this more realistic scenario, we augmented our model to include female preference loci, male trait loci, and loci at which introgressed alleles reduce viability. We found that the degree of purging of introgressed ancestry and the importance of mate choice’s bundling effect were similar to our baseline simulations (Fig. S4). This can be explained by the fact that most purging of introgressed DNA happens extremely quickly after admixture (Harris & Nielsen 2016, Veller et al. 2021), before the preference, trait, and ancestry loci have a chance to become fully dissociated by recombination.

We have considered a particular model of assortative mating in which a female prefers to mate with males of the species matching her majority ancestry. In an alternative model, a female most prefers males with the same ancestry as her (Irwin 2020), which might occur, for example, when mating is based on matching a polygenic trait like body size (Conte & Schluter 2013). Under this model of assortative mating (Methods), we find that the relative importance of the bundling effect is even greater than under our baseline “preference for conspecifics” model (Fig. S5B). This is because, in a model where hybrid females prefer hybrid males, the ancestry covariance among mating pairs (and therefore the trans-LD in offspring) is especially large. However, despite the increased importance of the bundling effect in this model, introgressed ancestry is not purged at an especially high rate (Fig. S5B), because hybrid males are not of especially low fitness—being favoured in mating by hybrid females (Irwin 2020). Similar results are observed for “sexual imprinting” models in which a female prefers to mate with males who have similar ancestry fractions to her father (Fig. S5C) or mother (Fig. S5D) (Irwin & Price 1999, Yang et al. 2019).

Thus far, we have considered only additive viability selection against introgressed ancestry. An alternative possibility is that the deleterious viability effects of introgressed alleles are largely recessive (Harris & Nielsen 2016). In that case, we might predict the bundling effect of mate choice—and in particular its trans channel—to have an especially large influence on the rate of purging of introgressed ancestry, as it generates an excess of homozygosity. In fact, in simulations of this scenario, we observe only a modest increase in the importance of the bundling effect (Fig. S6) relative to the additive case. The reason is that the overall influence of the trans channel is dominated by its effect on the *L*(*L* − 1) ≈ *L*^2^ possible trans associations across locus pairs, rather than its effect on the *L* possible within-locus trans associations (for which dominance is relevant).

A further possibility is that introgressed alleles reduce viability because of deleterious epistatic interactions with recipient-species alleles (Dobzhansky 1937, Muller 1942). Altering our model to include a large number of such incompatibilities, we find that the bundling effect of mate choice again has an important effect in enhancing viability and sexual selection (Fig. S7). It has been argued that persistent introgression of donor-species alleles should often cause the recipient-species alleles with which they are incompatible to be eliminated by selection, collapsing the post-zygotic barrier to gene flow (Barton & Bengtsson 1986, Bank et al. 2012). By accelerating the wholesale purging of introgressed alleles, the bundling effect dampens their impact on the frequencies of recipient-species incompatible alleles, promoting the maintenance of incompatibility-based post-zygotic barriers and thus species boundaries.

The strength of ancestry-based assortative mating can be quantified by the correlation coefficient of ancestry proportion between mates, *ρ*. In the Methods, we show that, if the ancestry correlation among mating pairs is *ρ*, then the bundling effect causes a factor 1 + *ρ* more introgressed ancestry to be purged in the generation immediately following assortative mating, and a factor ∼1/(1 − *ρ*) more introgressed ancestry to be purged in the long run. The latter calculation relies on some potentially unrealistic assumptions, notably that selection does not alter ancestry variance appreciably within each generation, but we may conservatively treat 1 + *ρ* and 1/(1 − *ρ*) respectively as lower and upper bounds for the overall impact of the bundling effect.

The most direct way to measure *ρ* is to estimate the ancestry proportions of mates. This requires detailed knowledge of mating pairs. One system where this is possible is the long-term study of baboons in Kenya’s Amboseli basin (Alberts & Altmann 2001). There, yellow baboons (*Papio cynocephalus*) and anubis baboons (*P. anubis*) hybridize (Alberts & Altmann 2001, Charpentier et al. 2012), and genomic analyses indicate that the minor anubis ancestry has been purged over time (Vilgalys et al. 2021). Tung et al. (2012) used long-term observations of mating behaviour in this system to investigate the determinants of mating success and mate pair composition, revealing ancestry-based assortative mating. Using data from Tung et al. (2012), we calculate an ancestry correlation of *ρ* = 0.195 among putative mating pairs (Methods; Fig. S8A). If anubis ancestry is deleterious, this estimate translates to a ∼20-24% increase in the purging of anubis ancestry due to the bundling effect.

Even in cases where mating is not observed, *ρ* can still be measured by estimating the ancestry fractions of mothers and offspring. In a hybrid population of the swordtail fish species *Xiphophorus birchmanni* and *X. cortezi* in Hidalgo, Mexico, genomic evidence suggests that the minor-parent ancestry (*birchmanni*) is deleterious to individuals of predominantly major-parent ancestry (*cortezi*) (Langdon et al. 2021). Powell et al. (2021) measured mother-offspring ancestry differences to infer paternal ancestry, finding evidence for strong ancestry-based assortative mating in this system [consistent with other swordtail systems (Culumber et al. 2014, Schumer et al. 2017)]. Using the mother-offspring data of Powell et al. (2021), we calculate an ancestry correlation among mating pairs of *ρ* = 0.928 (Methods; Fig. S8B). This estimate is consistent with a ∼2-14 fold increase in the purging of introgressed ancestry, revealing that in systems with strong but imperfect premating isolation, any ancestry that does introgress across the pre-zygotic barrier will experience much more rapid post-zygotic purging because of the bundling effect of mate choice.

Finally, to obtain a broad sense of the quantitative importance of the bundling effect, we turn to a meta-analysis of assortative mating in avian hybrid zones, which found an average correlation coefficient between mates of 0.44 (Randler 2008, Irwin 2020, Methods). Substituting this value into our calculations above, we estimate that the bundling effect of mate choice increases the purging of introgressed DNA by 44-80%.

Together, these estimates suggest that the bundling effect of mate choice plays an important role in the genetic isolation of species.

## Acknowledgments

We are grateful to Nate Edelman and members of the Coop lab at UC Davis for helpful discussions, and to Molly Schumer, Dan Powell, Jenny Tung, and Arielle Fogel for providing data and for help with data analysis. PM is supported by a Center for Population Biology postdoctoral fellowship and an NSF postdoctoral research fellowship. CV is supported by a Branco Weiss fellowship. This work was supported in part by the National Institute of General Medical Sciences of the National Institutes of Health (grants NIH R01 GM108779 and R35 GM136290 to G. Coop).

## Methods

### Admixture pulse

The model organism is a diploid sexual with a genome of length *L* = 1,000 loci. In our simulations, the “recipient species” population is of size *N* = 100,000, and experiences a sudden pulse of introgression such that a fraction *I*_*0*_ = 0.2 of the generation-0 population are of 100% donor species ancestry. Using the SLiM 3 simulation software (Haller & Messer 2019), we track the overall introgressed fraction in subsequent generations, *I*_*t*_.

### Viability selection

Introgressed alleles across the genome reduce viability in the recipient species, with the deleterious effect equal and additive across and within loci. Thus, if a fraction *I* of an individual’s diploid genome is introgressed, the individual’s viability is 1 − *IS*, where *S* is the strength of selection. In our simulations, *S* = 0.1.

### Mate choice

In our initial model, females engage in mate choice based on ancestry. All the models of mate choice that we consider are fixed relative preference models (Seger 1985): each adult female has a strength of preference for every adult male in the population, and chooses to mate with a given male with probability proportional to her strength of preference for him. The expected number of matings is the same for each adult female, so that only viability selection operates among females. In contrast, some males have a higher expected number of matings than others, so that both viability and sexual selection operate in males. Our baseline “preference for conspecifics” model for the strength of a female’s mating preference is

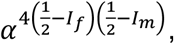

where is *I*_*f*_ is the female’s introgressed fraction, *I*_*m*_ is the male’s introgressed fraction, and *α* is the overall strength of mating preferences in the population. In the simulations displayed in the Main Text, *α* = 4; we explore various values of *α* in Fig. S2.

The first alternative assortative mating model that we consider (Fig. S5B) is

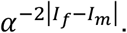

The “sexual imprinting” models that we consider (Fig. S5C,D) are

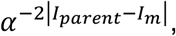

where *I*_*parent*_ is the introgressed fraction of the female’s father (Fig. S5C) or mother (Fig. S5D).

### The rate of purging is proportional to the population ancestry variance

Treating the ancestry proportion of an individual as a phenotype, the change in mean ancestry (*I*_*t*_) from one generation to the next can be written, using the breeder’s equation, as

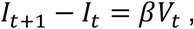

where *V*_*t*_ is the variance in ancestry proportion across individuals, as the ancestry proportion is a perfectly additive phenotype, and *β* is the directional selection gradient on ancestry. Under our additive model of viability selection against introgression, with no mate choice, 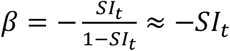 (Veller et al. 2021), and so the decrease in introgressed ancestry is proportional to the variance in ancestry proportion. Under more complex models of selection, e.g., viability selection due to Dobzhansky-Muller incompatibilities and sexual selection due to ancestry-based mate choice, our selection gradient would no longer take this simple form. However, the selection gradient can always be empirically calculated as the slope of fitness regressed on individual ancestry proportion; it will then naturally include fitness components due to mate choice and may depend on the ancestry composition of the population. The breeder’s equation will always predict the change in mean ancestry in the next generation as the product of the selection gradient and the population variance in ancestry. Thus, in a generation, we can always conceptualize the effect of mate choice in terms of its effect on ancestry variance and its effect on the strength of directional selection.

### Decomposition of the population variance in ancestry

An individual’s introgressed fraction *I* can be decomposed into maternal (*m*) and paternal (*p*) contributions:

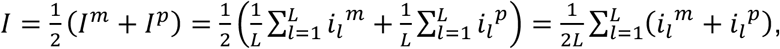

where *I*^*m*^ and *I*^*p*^ are the introgressed fractions of, respectively, the maternally and paternally inherited genomes of the individual, and *i*_*l*_^*m*^ and *i*_*l*_^*p*^ are indicator variables for whether the maternally and paternally inherited alleles at locus l are introgressed.

The ancestry variance across all individuals is

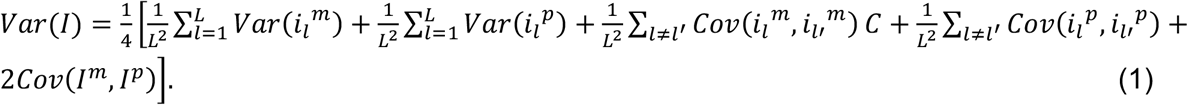

The term *Cov*(*I*^*m*^, *I*^*p*^) in Eq. (1) is the ancestry covariance between maternally and paternally inherited genomes, i.e., the overall trans-linkage disequilibrium (trans-LD). The terms 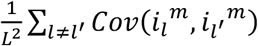 and 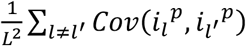 in Eq. (1) are the ancestry covariances within maternal and paternal genomes, i.e., the cis-linkage disequilibrium (cis-LD). In the absence of trans-LD and cis-LD, ancestry variance would simply be a function of the allele frequencies at different loci: 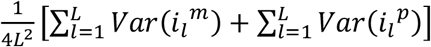 in Eq. (1).

The variance, trans-LD, and cis-LD values displayed in Fig. 3 were calculated as above, and normalized by dividing through by a factor of *I*_*t*_ (1 – *I*_*t*_), where *I*_*t*_ = *E*[*I*] is the population’s introgressed fraction in generation *t*. This normalization accounts for the fact that the variances and covariances scale with the overall frequency of introgressed ancestry; under this normalization, the overall variance, trans-LD, and cis-LD are the same for the “no mate choice” and “mate choice with bundling removed” cases in Fig. 3, despite introgressed ancestry being purged at a higher rate in the latter case owing to the additional effect of sexual selection.

### Sex chromosomes

In the configurations of our model that involve sex chromosomes, the ancestry fraction of a heterogametic individual is calculated as *I* = (*AI*^*A*^ + *XI*^*X*^)/(*A* + *X*), where *A* and *X* are the autosomal and X-linked (or Z-linked) fractions of total haploid genome length, respectively, and *I*^*A*^ and *I*^*X*^ are the individual’s autosomal and X-linked introgressed fractions. Notice that, in treating the hemizygous X equivalently to the autosomes in this calculation, we are assuming full dosage compensation. For the stylized genomes we consider in Fig. S3, *A* = *X* = 1/2.

### Recombination maps

Loci were assumed to be spaced evenly along the physical (bp) genome. For stylized recombination processes, we assumed that all loci were unlinked (e.g., Figs. 2A, 3) or that the rate of recombination between adjacent locus pairs on the same chromosome was constant (Fig. S3). In the case of realistic recombination processes (Fig. 2B), we interpolated empirical linkage maps along our evenly spaced loci. For humans, we used the male and female maps generated by Kong et al. (2010); for *D. melanogaster*, we used the female linkage map produced by Comeron et al. (2012). We ignored crossover interference in our simulations.

### Dobzhansky-Muller incompatibilities

We considered only pairwise Dobzhansky-Muller incompatibilities (DMIs) between donor and recipient species alleles (i.e., no higher-order epistasis). Our simulations involved *L* = 1,000 loci, harboring *D* = 100 non-overlapping DMI locus pairs, and 800 “ancestry loci”.

Suppose that the incompatible alleles at loci *l*_1_ and *l*_2_ are A and B, with a and b being the alternative alleles at these loci, respectively. We considered two dominance cases for our DMIs (Turelli & Orr 2000). In the first case, all DMIs are of intermediate epistatic dominance, so that the genotypes AaBb, AABb, AaBB, and AABB suffer viability reductions *s*/2, 3*s*/4, 3*s*/4, and *s*. In the second case, all DMIs are epistatically fully recessive, so that only the genotype AABB suffers a viability reduction, of relative size *s*. In our simulations, *s* = 0.02, with fitness effects combining multiplicatively across DMIs. While viability selection in this model is based on epistasis across the *D* DMI locus pairs, mate choice is based on ancestry at all *L* loci.

If introgressed ancestry were to start at a low frequency, it would be strongly disfavored in mate choice, and sexual selection based on overall ancestry would overwhelm viability selection based on DMIs. In this case, the trajectories would resemble those for additive viability selection, as considered elsewhere in this paper. Therefore, to better isolate the role of DMIs themselves, we begin our simulations with a high fraction of introgressed ancestry (47.5%). In this case, sexual selection against introgressed ancestry is initially weak and most selection is due to DMIs. We track the overall fraction of introgressed ancestry through time, as well as the fraction of introgressed alleles at loci where the introgressed allele is involved in an incompatibility.

### Separate preference, trait, and ancestry marker loci

In the simulations that distinguished the loci underlying the female preference, the sexually selected male trait, and ancestry/viability fitness, we considered two architectures: (i) *P* = 100 preference loci, *T* = 100 trait loci, and *A* = 800 ancestry loci; (ii) *P* = 10 preference loci, *T* = 10 trait loci, and *A* = 980 ancestry loci. The strength of a female’s preference for a male was calculated as

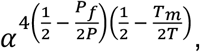

where *P*_*f*_ is number of introgressed preference alleles carried by the female and *T*_*m*_ is the number of introgressed trait alleles carried by the male. Both viability fitness and the overall introgressed fraction were determined according to the fraction of introgressed alleles at ancestry loci.

### Relating ancestry correlations between mates to increased population variance

Since there are many more locus pairs than individual loci (∼*L*^2^ vs. *L*) and introgressed alleles initially appear in perfect cis-LD with one another, the initial population ancestry variance, *V*_*0*_, is almost entirely tied up in cis-LD between introgressed alleles, with heterozygosity at the *L* individual loci in the genome contributing negligibly. For the same reason, the ancestry variance in generation *t* can be written as a sum of the total cis-LD and the total trans-LD:

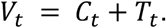

We will assume that all loci are unlinked (*r* = 1/2), which, for our purposes (short timescales), is approximately the case for most species (Crow 1988, Veller et al. 2019). In the construction of generation *t* + 1, recombination reduces the cis-LD from generation *t* by a factor of 1/2 and converts the trans-LD in generation *t* to new cis-LD at rate 1/2. If assortative mating among generation-t parents generates an ancestry correlation between mates of *ρ*, then an amount *ρV*_*t*_/2 of trans-LD is present in generation *t* + 1 (e.g., Zaitlen et al. 2017). Therefore,

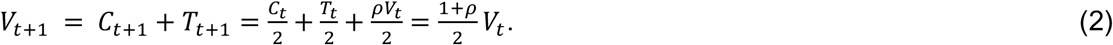

The amount of introgressed DNA purged in generation *t* is

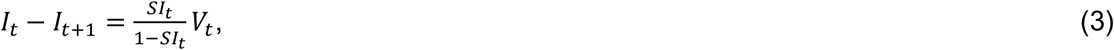

where *I*_*t*_ is the introgressed fraction in generation *t* and *S* is the overall strength of selection against introgressed ancestry (Veller et al. 2021). In the absence of assortative mating (*ρ* = 0), 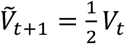. The amount of introgressed DNA purged in generation *t* + 1, given assortative mating in generation t, is 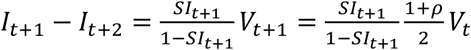, while the amount that would have been purged in the absence of the bundling effect of mate choice (same *S* but *ρ* = 0) is 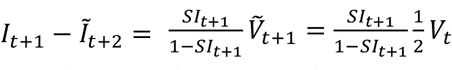 Therefore, the additional amount of introgressed DNA that is purged in generation *t* + 1 because of ancestry bundling induced by assortative mating in generation *t* is

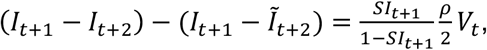

so that the bundling effect has increased purging by a factor of

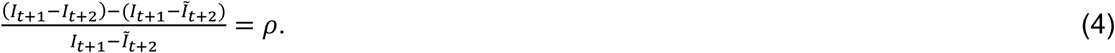

To understand how such effects compound over generations, we assume that natural and sexual selection are weak, such that nearly all dissipation of variance is due to recombination rather than selection. We further assume that the ancestry correlation within mating pairs is a constant value *ρ* each generation. Under these assumptions, we may iterate Eq. (2), yielding

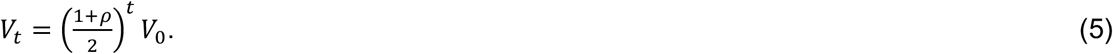

The amount of introgressed DNA purged in generation t is

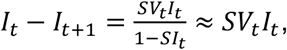

since *S* is assumed to be small. The total proportion of introgressed ancestry purged up to generation *t* can therefore be written

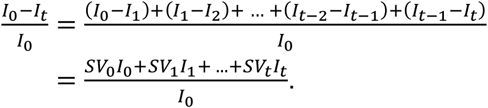

Since selection is assumed to be weak, *I*_*t*_ changes slowly, so that

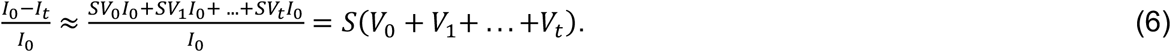

In the presence of mate choice, we substitute (5) into (6) to find

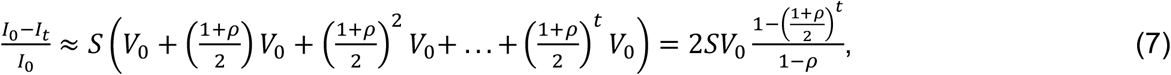

while, in the absence of the bundling effect of mate choice (same *S* but *ρ* = 0), the total proportion of introgressed ancestry purged up to generation *t* would instead be

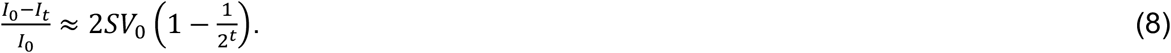

The excess fraction of purging due to the bundling effect of mate choice is therefore given by [Eq.(7) – Eq.(8)] / Eq.(8):

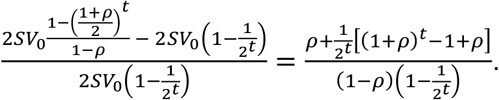

As *t* grows large, and assuming *ρ* < 1, this expression converges to *ρ*/(1 − *ρ*). Thus, eventually, overall purging has been increased by a factor of

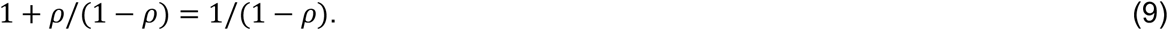

When selection is strong, ancestry variance will decay not only because of recombination, but because of selection as well; this will act to diminish the effect of assortative mating in decelerating the decay of ancestry variance.

### Empirical examples

#### Hybrid zone between yellow and anubis baboons

In this hybrid zone, rank and other social factors have been shown to play an important role in mate choice (Tung et al. 2012). In addition, Tung et al. (2012) found that males with more anubis ancestry are favoured in mate choice overall, and that there is also ancestry-based assortative mating. If, as genomic evidence suggests (Vilgalys et al. 2021), the minor parent ancestry (anubis) has historically been deleterious in this population, then our calculations above imply that it is the ancestry correlation coefficient within mating pairs that determines the impact of ancestry bundling on the purging of anubis ancestry. In particular, while there is a directional mating advantage of anubis ancestry, this is part of overall direct selection on ancestry, and therefore included in the selection gradient along with other factors, while assortative mating increases ancestry variance (see section “The rate of purging is proportional to the population ancestry variance” above).

To calculate the correlation coefficient among putative mating pairs, we filtered the data of Tung et al. (2012) to include only those male-female pairs where consortship behaviour (mate guarding) was observed in a period when the female conceived—these are putative mating pairs. In cases where multiple males consorted the same female in a single conceptive period, we randomly selected one of the males, and then calculated the ancestry correlation coefficient among consorting male-female pairs. Repeating this male-sampling procedure 100,000 times, we calculated an average ancestry correlation coefficient within mating pairs of *ρ* = 0.195. This value is consistent with a 19.5% increase in purging of anubis ancestry in the generation after assortative mating [Eq. (4)], and a ∼24.2% increase in long-term purging [Eq. (9)].

#### Swordtail fish

In a hybrid population between *Xiphophorus birchmanni* and *X. cortezi*, Powell et al. (2021) measured genome-wide ancestry fractions in mothers and their embryos. We use these measurements to calculate the correlation coefficient between mating pairs, *ρ*. Let *M, F*, and *O* be minor-parent ancestry (*birchmanni*) fractions of a mother, father, and offspring respectively. Then

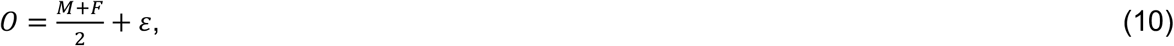

where *ε* is a noise term due to random segregation in the maternal and paternal meioses, with,*Cov*[*M*, ε] = 0. From Eq. (10),

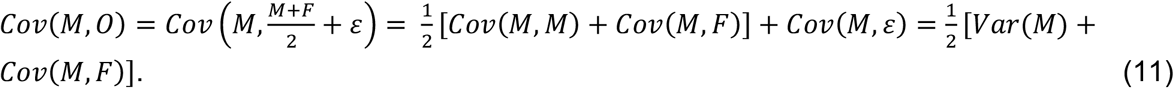

Using Eq. (11), the slope of the regression of offspring ancestry on maternal ancestry is

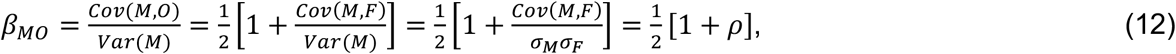

where *σ*_*M*_ and *σ*_*F*_ are the ancestry standard deviations of mothers and fathers, which we assume to be equal (such that *Var*(*M*) = *σ*_*M*_*σ*_*F*_). Rearranging Eq. (12),

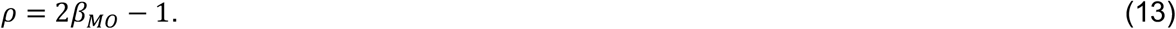

Since some mothers have multiple embryos in the data of Powell et al. (2021), we calculated *β*_*MO*_ using the average ancestry fraction of each mother’s offspring, yielding an estimate of *β*_*MO*_ = 0.964 (note that the same value would be obtained asymptotically if we averaged over iterations in which we randomly choose one embryo per mother and calculate *β*_*MO*_ for the reduced dataset). From Eq. (13), our estimate of *β*_*MO*_ = 0.964 corresponds to a correlation coefficient among mating pairs of *ρ* = 0.928. This value is consistent with a 1.928-fold increase in purging of minor parent ancestry in the generation after assortative mating [Eq. (4)], and a 13.80-fold increase in long-term purging [Eq. (9)].

#### Meta-analysis in birds

Randler (2008) carried out a meta-analysis of assortative mating in avian hybrid zones, and found an average *z*-score of 0.44, which, by Fisher’s *z*-transformation, corresponds to a correlation coefficient of *ρ* = 0.44 (Rosenberg et al. 2000, Irwin 2020). Randler’s (2008) meta-analysis covered cases of assortative mating based on species-diagnostic phenotypes and on genetic ancestry. If the calculated correlation coefficient applies to ancestry-based assortative mating, then it is consistent with a 44% increase in purging of minor parent ancestry in the generation after assortative mating [Eq. (4)], and a 79% increase in long-term purging [Eq. (9)].

## Supplementary figures

**Figure S1.**
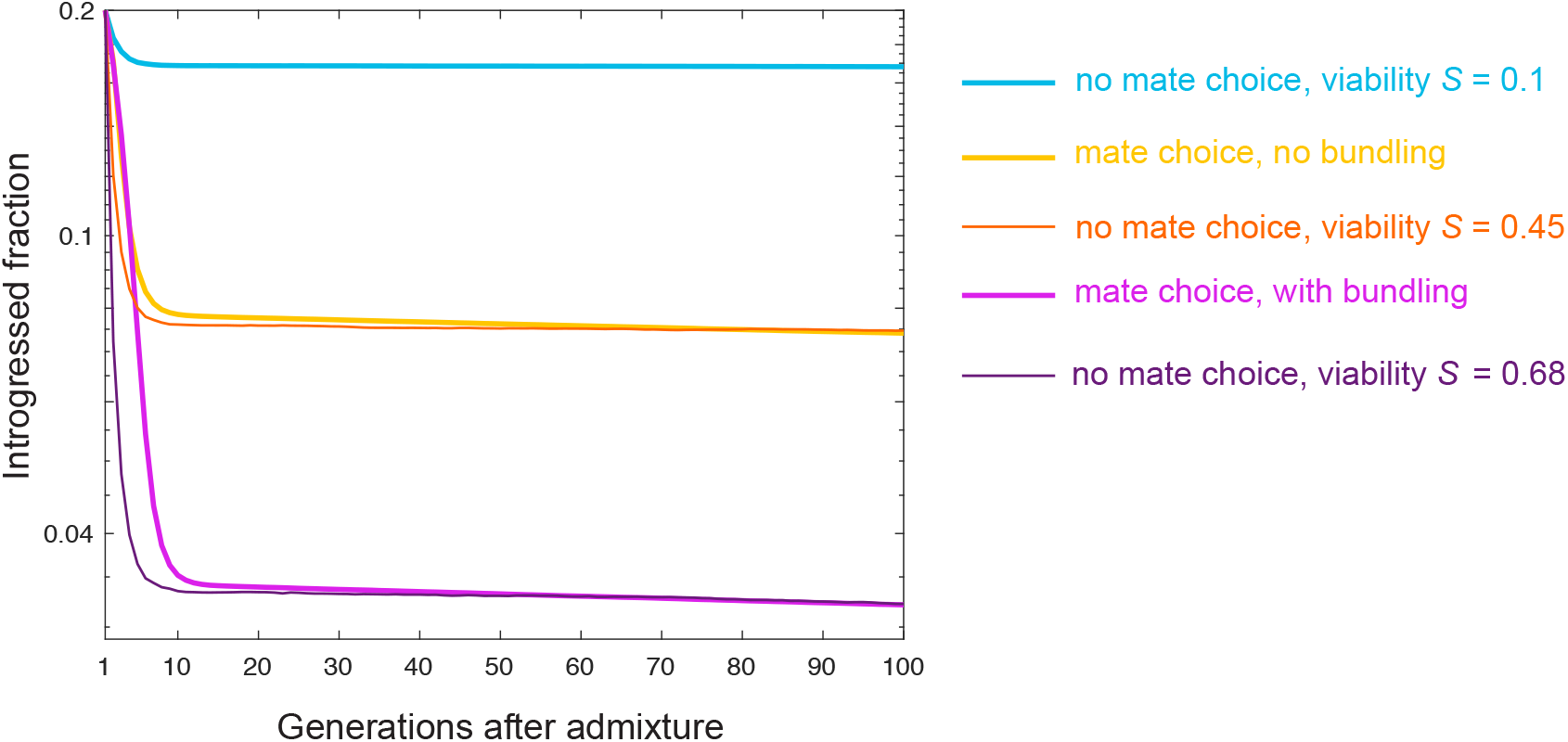
The bundling effect of mate choice enhances the effective strength of selection against introgressed ancestry. In our baseline simulations, in the absence of mate choice, viability selection acts against introgressed alleles with total effect *S* = 0.1. With mate choice, selection against introgressed ancestry is stronger, owing to the additional effects of sexual selection and bundling. To generate a common metric for the strength of these additional effects, we determined—by visual inspection—how strong viability selection alone would need to be to mimic the trajectories of the purging of introgressed ancestry under the two mate choice scenarios in our simulations: mate choice with its bundling effect artificially removed, and mate choice with its bundling effect left intact. Parameters are as in Fig. 2A.

**Figure S2.**
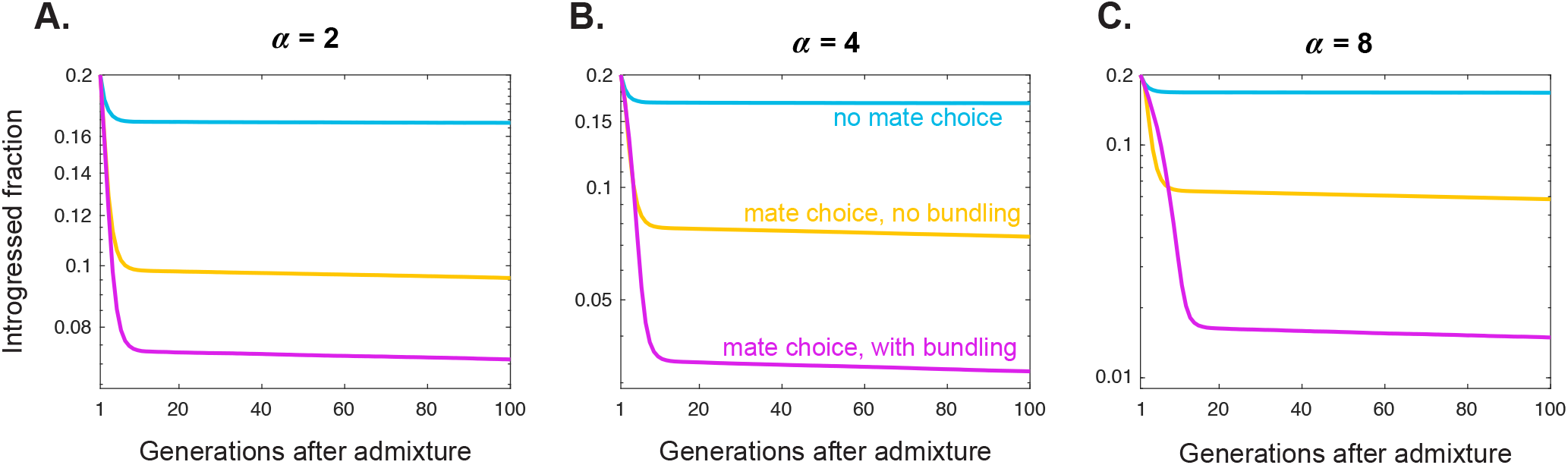
The importance of the bundling effect increases with the strength of assortative mating, *α*. Parameters in **B** are as in Fig. 2A; relative to this baseline case, mate choice is weaker in **A** and stronger in **C**.

**Figure S3.**
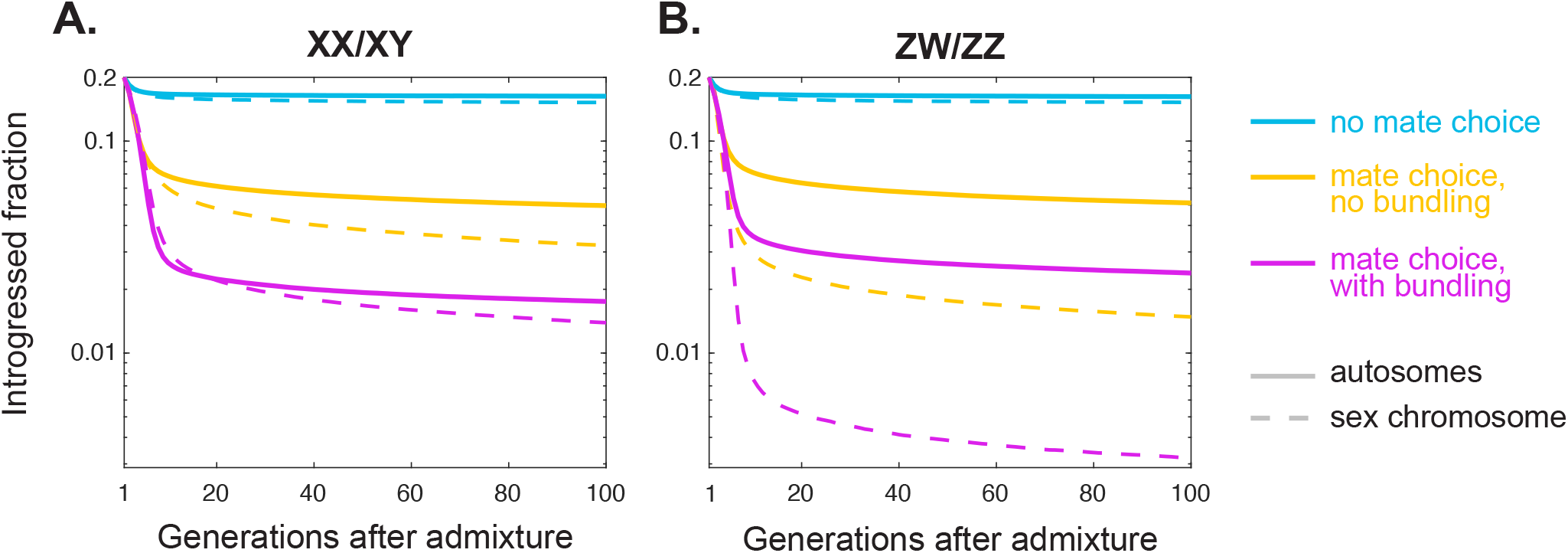
Rates of purging of introgressed ancestry on sex chromosomes and autosomes in male- and female-heterogametic systems. Under both male (**A)** and female (**B)** heterogamety, the sex chromosome purges more introgressed ancestry than the autosomes, in all three simulation setups: no mate choice, mate choice with its bundling effect removed, and full mate choice. The increase in the rate of sex-linked purging under mate choice, and the effect of bundling, are especially large in female-heterogametic systems (**B**). In these simulations, the genome comprises one sex chromosome and one autosome, each containing 500 loci. Recombination is uniform along the autosome and the sex chromosome, with an average of one crossover per chromosome per gamete. There is no recombination along the sex chromosome in the heterogametic sex. Otherwise, parameters as in Fig. 2A.

**Figure S4.**
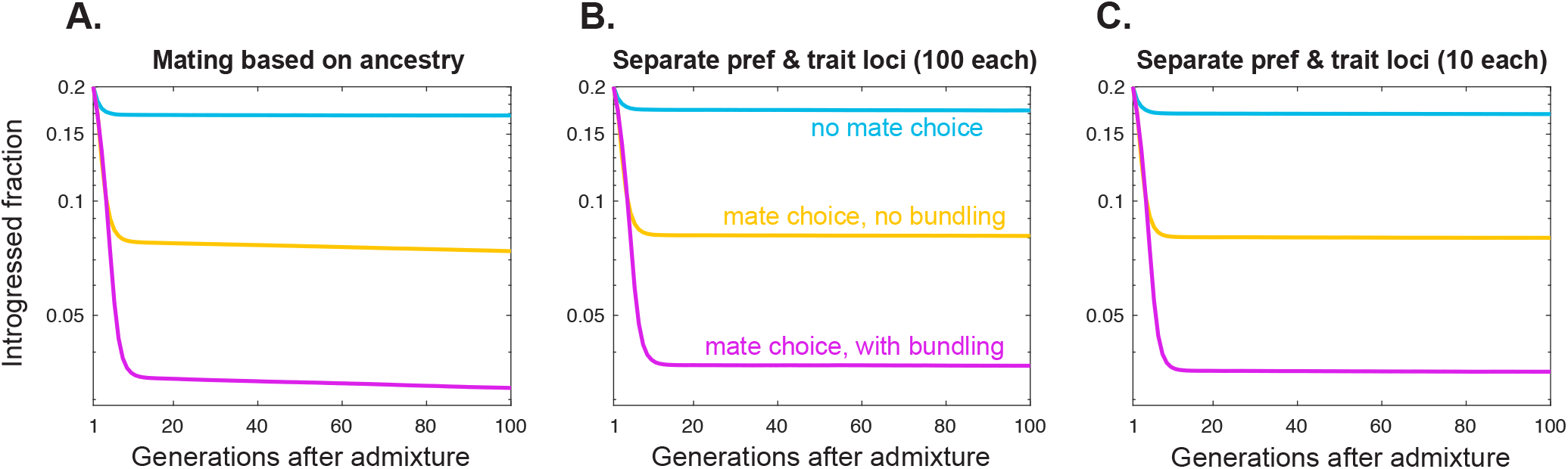
Purging of introgressed ancestry in a genetically more realistic model. **A**. Purging under our baseline model where viability and sexual selection depend only on genome-wide ancestry proportions. **B**,**C**. Purging in a model where a female mating preference, the sexually selected male trait, and viability fitness are each encoded at distinct sets of loci. In **B**, 100 loci underlie the female preference, 100 loci underlie the male trait, and 800 loci underlie viability fitness. In **C**, 10 loci underlie the female preference, 10 loci underlie the male trait, and 980 loci underlie viability fitness. The introgressed fractions displayed are calculated from the viability loci. The resulting trajectories are similar to those in **A**. Parameters as in Fig. 2A, and all loci are unlinked.

**Figure S5.**
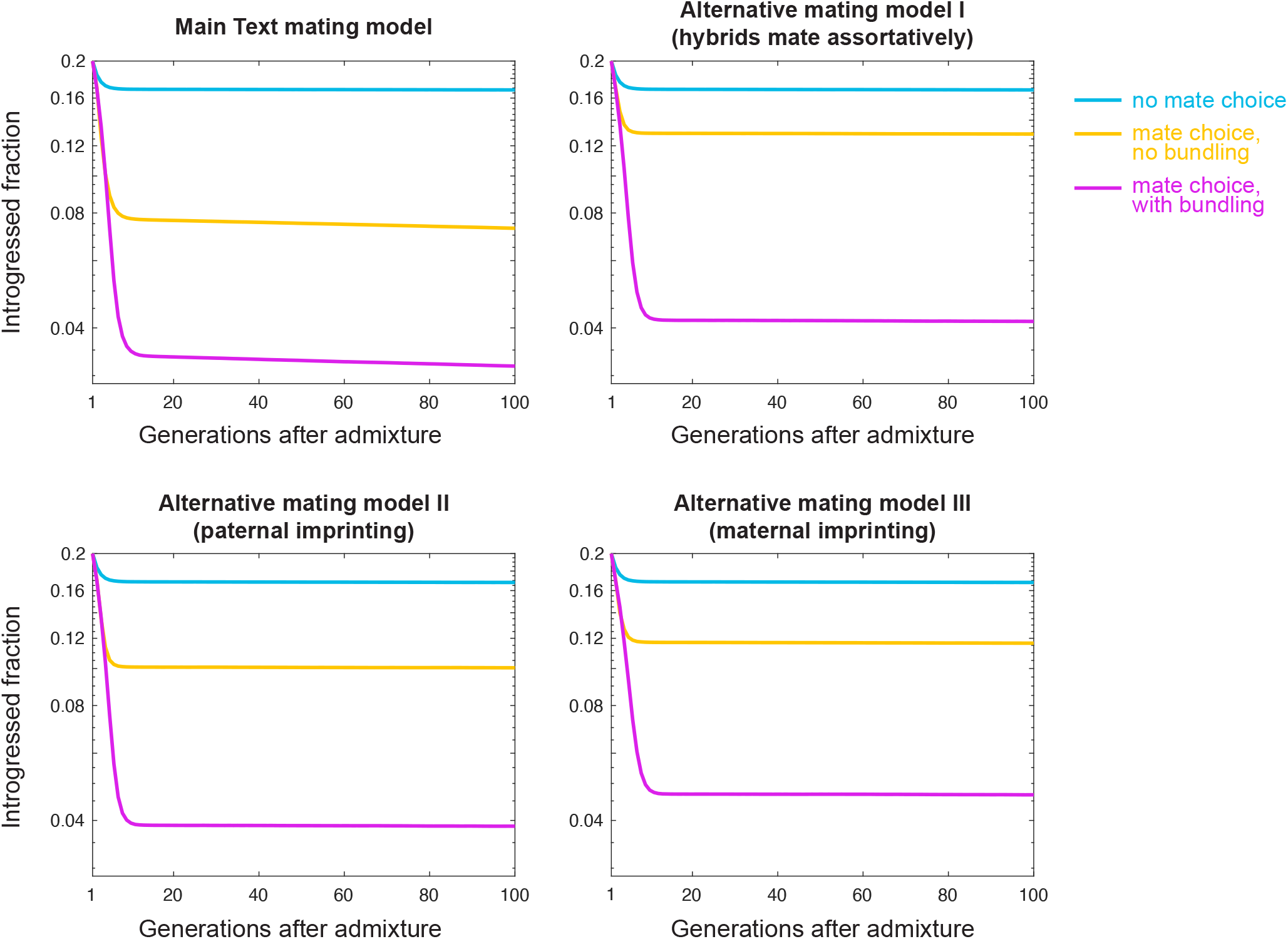
Purging under alternative specifications of assortative mating. **A**. Purging under our baseline model of assortative mating, in which females with majority species-X ancestry most prefer males of 100% species-X ancestry. **B**. Purging under a model of mate choice where females with a fraction *x* of species-X ancestry most prefer males who also have a fraction *x* of species-X ancestry (Methods). Under this alternative model, the ancestry correlation between mates is stronger, but the overall effect of mate choice on the purging of introgressed DNA is weaker because hybrid males can be favoured in mate choice. Therefore, the overall effect of mate choice on the purging of introgressed DNA is smaller than in our baseline model, but the proportion due to the bundling effect is larger. **C**,**D**. Purging under models in which a female whose father (**C**) or mother (**D**) has a fraction *x* of species-X ancestry most prefers males who also have a fraction *x* of species-X ancestry (Methods). In these “sexual imprinting” models, purging is slightly slower, and the bundling effect slightly weaker, than in **B**. Parameters as in Fig. 2A.

**Figure S6.**
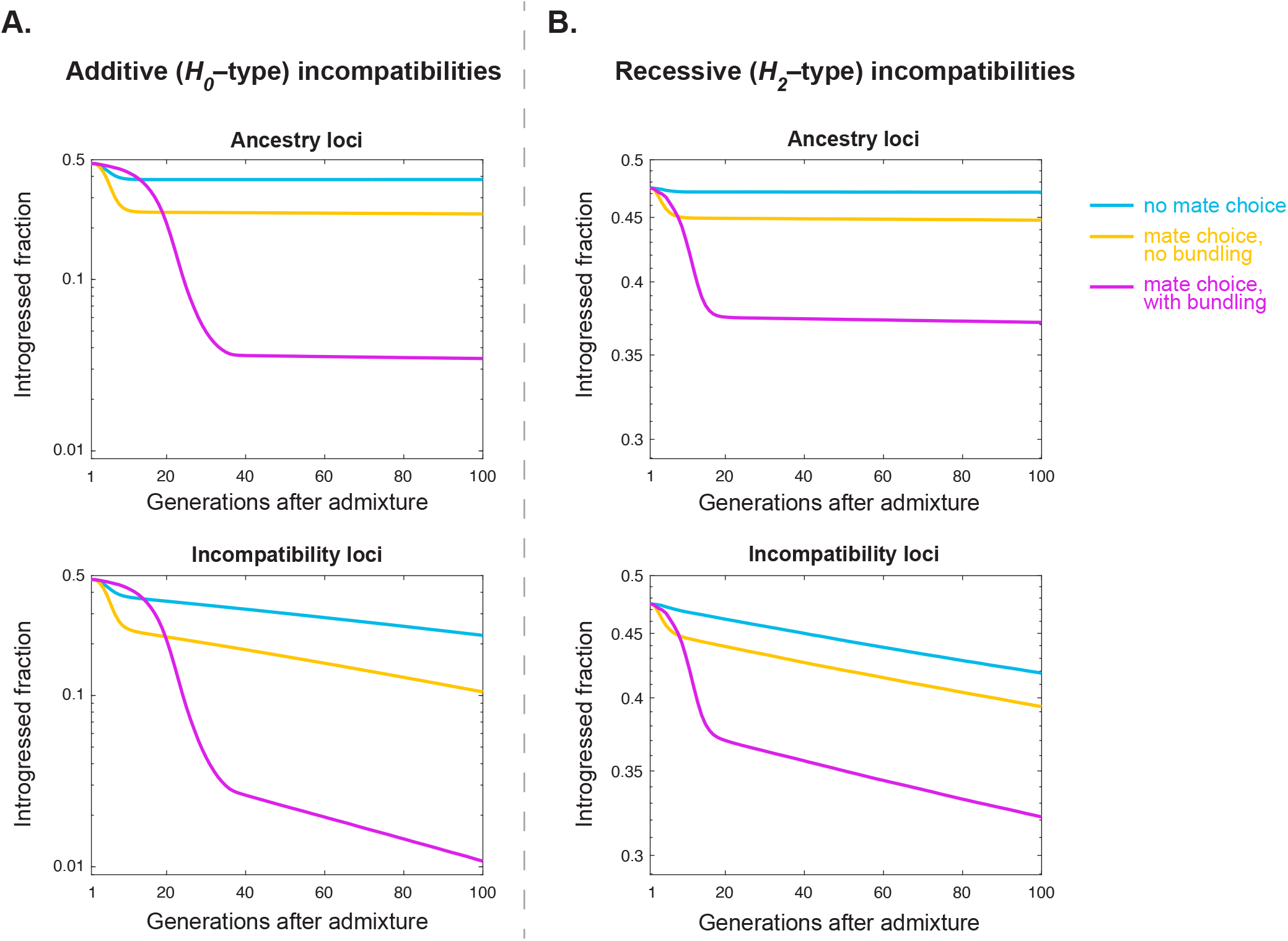
The bundling effect is important when viability selection against introgressed ancestry is due to Dobzhansky-Muller incompatibilities. **A**. The case where selection against incompatible alleles shows intermediate epistatic dominance. **B**. The case where selection against incompatible alleles is epistatically recessive. In both cases, we modelled 100 non-overlapping DMI locus pairs in a genome of 1,000 loci, with the maximum viability loss for each DMI being 2%, and viability effects combining multiplicatively across DMIs. Top panels track introgressed fractions at the 800 “ancestry” loci (i.e., non-DMI loci), while bottom panels track the introgressed fraction at the 100 loci at which the introgressed allele is involved in an incompatibility. Other parameters as in Fig. 2A. Note the difference in y-axis scales between **A** and **B**, to account for the fact that selection is weaker, and therefore less purging occurs, in the case where the incompatibilities are epistatically recessive.

**Figure S7.**
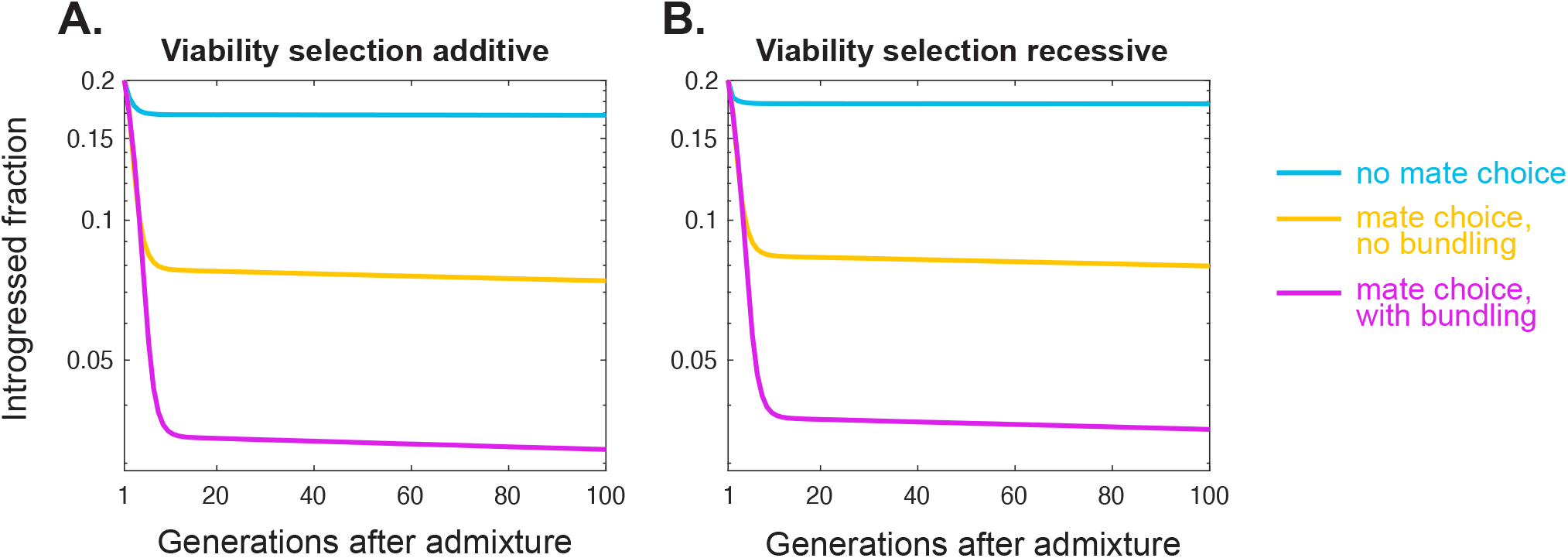
The bundling effect is not substantially more important when the deleterious effect of introgressed alleles on viability is recessive. **A**. Our baseline case, where viability selection against introgressed alleles is additive. **B**. The case where viability selection against the introgressed allele at each locus is recessive. Naively, one might expect the bundling effect to be especially important here, since it involves trans covariances that generate an excess of introgressed-allele homozygotes at each locus. However, this effect on the efficiency of viability selection is proportional to the number of loci, *L*, while the excess ancestry variance on which sexual selection acts is proportional to the number of locus pairs, ∼*L*^2^, explaining why the importance of the bundling effect is not much higher in this case, relative to **A**. Parameters as in Fig. 2A.

**Figure S8.**
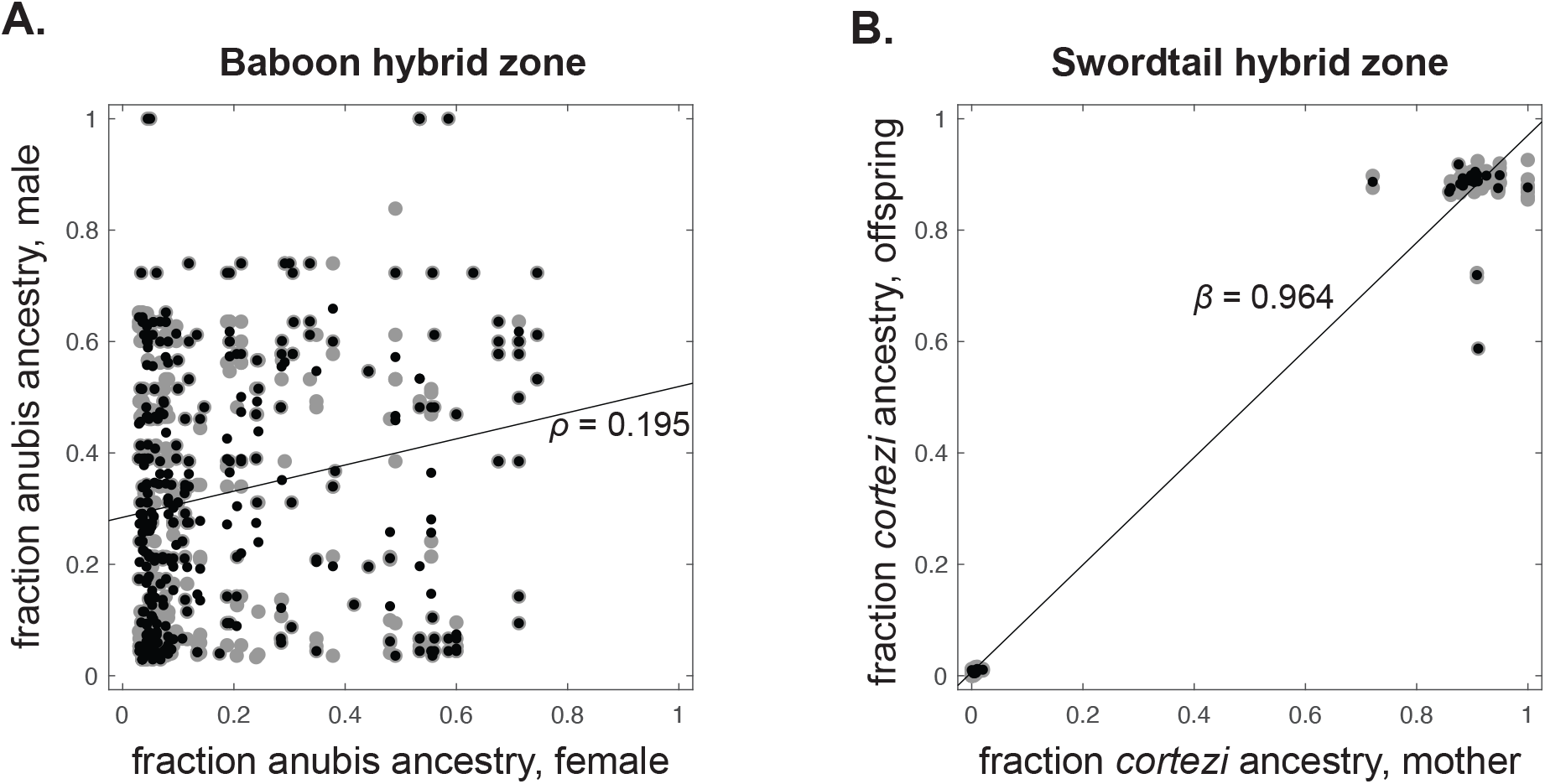
Assortative mating in two hybrid zones. **A**. Ancestries of females and their putative mates in a hybrid zone between yellow and anubis baboons. Grey dots display the ancestry of each female who conceived in a given period vs. each male who consorted with the female in that period; black dots show the average ancestry among all males who consorted with the female in that period. The displayed ancestry correlation coefficient between mating pairs, *ρ* = 0.195, is calculated via a bootstrap procedure described in the Methods. **B**. Ancestries of females and their offspring in a hybrid population of the swordtail species *Xiphophorus birchmanni* and *X. cortezi*. Grey dots represent each mother-offspring pair, while black dots display the average ancestry among each female’s offspring. The displayed regression slope, *β* = 0.964, is calculated using the average offspring values, and, as detailed in the Methods, translates to an ancestry correlation coefficient among mating pairs of *ρ* = 0.928.

